# Ageing impairs protein leveraging in a sex-specific manner in *Drosophila melanogaster*

**DOI:** 10.1101/2022.08.05.502931

**Authors:** Helen J. Rushby, Zane B. Andrews, Matthew D. W. Piper, Christen K. Mirth

**Affiliations:** School of Biological Sciences, Monash University, Melbourne, VIC 3800, Australia

**Keywords:** age-related decline, feeding behaviour, food intake, macronutrient balancing, protein leveraging, sex-specific behaviour

## Abstract

Modifying the relative proportions of macronutrients in an animal’s diet has noteworthy effects on its reproduction, lifelong health, and lifespan. Because of this, a wide range of animals carefully regulate their nutrient intake toward species and stage-specific targets. However, when animals are unable to reach their nutrient target from their existing food resources, they will compromise between overconsuming one nutrient and under-consuming the deficit nutrient. In this study, we used capillary feeding (CAFE) assays to understand the rules of compromise of adult fruit flies (*Drosophila melanogaster*) of different sex, mating status, and age when constrained to single diets. We found that young male and female *D. melanogaster* compromised by consuming more food on diets with low protein to carbohydrate (P:C) ratios compared to diets with high P:C ratios. Further, young male and female flies varied their carbohydrate intake significantly more than their protein intake, and female flies varied their carbohydrate intake significantly more than males. To test for effects of mating status on nutrient intake, we compared food intake of young mated and virgin females. We found that both virgin and mated females compromised by consuming more food on the low P:C diet compared to high P:C diets; however, mated females consumed more food than virgin females. As flies aged, they decreased their overall food intake and showed more modest alterations in their food intake across varying P:C diets. Further, mated females ceased to compromise for the protein deficit at a younger age than males. These findings provide new understanding about differences in protein leveraging behaviour across sexes, and how these behaviours change with age.

**Highlights:**

- Young fruit flies exhibit protein leveraging behaviour, varying their carbohydrate consumption more than protein
- Young mated female flies vary their carbohydrate consumption significantly more than young males
- Both virgin and mated female flies balance their nutrient intake similarly
- As flies age, their ability to protein leverage declines, and this occurs faster in female flies

## INTRODUCTION

Animals must ingest adequate amounts of high-quality nutrients to obtain energy and building blocks for homeostasis and physical activity. These energy resources are obtained from three macronutrients - protein, carbohydrates, and lipids. Altering the relative proportions of macronutrients in a diet has a considerable impact on reproduction, lifelong health, and lifespan across a broad range of taxa (Simpson & Raubenheimer, 2012). Since ingesting a diet with the optimal ratio of nutrients maximises an animal’s fitness and health, animals balance their macronutrient intake toward certain targets. Macronutrient balancing by altering feeding behaviour is reported widely in literature and across a variety of taxa (Raubenheimer & Simpson, 1997; Simpson et al., 2015). However, it is not yet known if this behaviour is modified by sex or mating status, and to what extent age alters food intake.

Animals forage selectively for nutrients that optimise certain life-history outcomes (Simpson et al., 2004). For instance, *Drosophila melanogaster* larvae target their nutritional intake towards protein to carbohydrate (P:C) ratios that minimize development time (Rodrigues et al., 2015). Adult female *Drosophila*, predatory beetles, and crickets regulate their food intake towards P:C ratios that maximize lifetime egg production (Bowman & Tatar, 2016; Camus et al., 2018; Jensen et al., 2012; Lee et al., 2008; Maklakov et al., 2008; Ribeiro & Dickson, 2010; Tsukamoto et al., 2014). Additionally, altering the ratio of P:C in a diet has profound effects on lifespan and lifelong health. For example, the lifespan of many insects and mice can be extended by decreasing the P:C ratio of a diet below the level that optimises reproduction (Le Couteur et al., 2016). As such, animals tune their foraging behaviour to achieve nutrient intake targets that optimise traits that are important for growth and reproduction.

When animals are constrained to a diet which does not allow them to reach their intake target, they will compromise between over-consuming one nutrient and under-consuming the deficit nutrient; this is known as the “rules of compromise” (Raubenheimer & Simpson, 1997). Both mice and humans tend to prioritise a constant protein intake, a behaviour known as protein leveraging (Simpson & Raubenheimer, 2005). Specifically, when organisms consume a diet that is disproportionately low in protein, they will protein leverage by eating more food as they try to ingest a target amount of protein (Felton et al., 2009; Gosby et al., 2011; Simpson et al., 2003; Simpson & Raubenheimer, 2005; Sørensen et al., 2008). Since changing the balance of macronutrients in a diet can result in overconsumption of energy, eating imbalanced diets in which carbohydrate or fat dilutes protein may be an important factor contributing to the current obesity epidemic (Simpson et al., 2015). Furthermore, animals maintained on a high protein, low carbohydrate diet have increased risk for conditions associated with overeating, such as high blood pressure, diabetes, and heart disease (Floegel & Pischon, 2012; Lagiou et al., 2012; Solon-Biet et al., 2014). Hence, the regulation of nutrient intake is exceptionally important to lifelong health.

Although overconsumption is associated with several harmful conditions, some individuals may benefit from increased food intake. For example, there is a high prevalence of malnutrition among the elderly (Shahar et al., 2002). The majority of this population experiences suppressed appetite, a decrease in appreciation for food flavour, and a greater reduction in appetite after high protein meals compared to young and middle-aged individuals (Kremer et al., 2007; Morley, 1997; Paddon-Jones & Leidy, 2014). Further, elderly individuals struggle to return to their previous weight after weight loss, suggesting an impaired ability to regulate food intake (Roberts, 2000). Given the high occurrence of malnutrition and the impaired regulation of food intake in the elderly, the ability to regulate intake targets effectively may change with age. To understand how this occurs, experiments conducted across organisms’ lifespans are required to disentangle the effects of ageing on the rules of compromise.

The fruit fly, *Drosophila melanogaster*, provides an excellent model for exploring the effects of age, sex, and reproductive status on rules of compromise behaviour. This is because it is easy to rear large numbers of individuals in age-matched cohorts, and there are well-established methods to generate virgin and mated female flies. The short lifespan of the fruit fly, relative to vertebrates, allows for relatively quick longitudinal studies where we can assess the effects of age on macronutrient balancing across adult lifetime. Thus far, it is known that *D. melanogaster* larvae exhibit protein leveraging behaviour, consuming more food overall to compensate for a diet low in protein relative to carbohydrate (Carvalho & Mirth, 2017). However, larvae have different nutritional requirements than adults due to their need to support growth, which requires higher concentrations of protein (Rodrigues et al., 2015). Therefore, this study investigated the rules of compromise used by the adult fly and how their rules are affected by sex, mating status, and age.

To quantify rules of compromise, we used Capillary Feeding (CAFE) assays (Ja et al., 2007) to constrain *D. melanogaster* flies of different sex, age, and mating status to single diets that ranged in their P:C composition. Since mating status and sex have already been shown to alter nutrient choices (Bowman & Tatar, 2016; Camus et al., 2018; Lee et al., 2013; Maklakov et al., 2008; Ribeiro & Dickson, 2010; Tsukamoto et al., 2014), we hypothesized that the rules of compromise could differ across sexes, with mating status in females, and with age. Specifically, we predicted that females would show a stronger priority towards protein intake than males and that this effect would be greater post-copulation. Additionally, we expected that the flies’ ability to make compromises for a nutrient deficit would break down as they age, as the anabolic burden for reproduction falls and ageing-related physiological decline sets in. This study will lay the foundation for studies of potential mechanisms underlying changes in feeding behaviours across age groups.

## METHODS

### Fly stocks and maintenance

All experiments were conducted with a laboratory stock of outbred wild type *D. melanogaster* flies (Red Dahomey). Flies were reared by performing 24-hour egg lays as described in Linford et al. (2013). Eggs were collected and washed in phosphate buffered saline and 25 µL of eggs were pipetted into 70 mL of sucrose/yeast (SY) media in 250 mL bottles. SY media contained 10 g of Grade J3 agar (Gelita Australia Pty Ltd.; order code: A-181017, 100 g of autolysed Brewer’s Yeast (MP Biomedicals; order code: 903312; ∼45% protein, ∼45% amino acids, ∼1% fat, ∼5% moisture, ∼4% other), and 50 g of sucrose (Bundaberg Australia; order code: M180919) in 1 L of water. We added 30 mL of nipagin (Sigma-Aldrich; order code: W271004-5KG-K) in absolute ethanol (Thermo-Fischer; order code AJA214-2.5LPL) and 3 mL of propionic acid (Merck, Pty Ltd., Germany; order code: 8.00605.0500) as preservatives to prevent microbial growth.

Flies were maintained at 25°C and 60-80% humidity with a 12h:12h light:dark cycle. For CAFE assays, liquid experimental media comprised sucrose, autolysed Brewer’s yeast, and water. The amount of yeast relative to the amount of sucrose in water was altered to achieve ratios of 1:1 P:C, 1:4 P:C, and 1:16 P:C for low (1038 KJ/L) and high (2071 KJ/L) calorie diets (**Table 1**). 1% blue food dye (Queen) was added for contrast. The food was autoclaved to reduce spoilage.

**Table 1.** High and low calorie 1:1, 1:4 and 1:16 P:C diets were produced by altering the amount (grams) of sucrose and yeast in 1 L of water

|  | High calorie<br>(2071 KJ/L) | Low calorie<br>(1038 KJ/L) |
| --- | --- | --- |
| 1:1 P:C |  |  |
| Yeast | 142.22 | 71.11 |
| Sucrose | 17.07 | 8.54 |
| Protein | 64.00 | 32.00 |
| Carbohydrates | 64.00 | 32.00 |
| 1:4 P:C |  |  |
| Yeast | 56.89 | 28.45 |
| Sucrose | 83.63 | 41.82 |
| Protein | 25.61 | 12.80 |
| Carbohydrates | 102.40 | 51.20 |
| 1:16<br>P:C |  |  |
| Yeast | 16.73 | 8.37 |
| Sucrose | 115.00 | 57.5 |
| Protein | 7.53 | 3.76 |
| Carbohydrates | 120.52 | 60.26 |

### Fly rearing and CAFE assay set up

Flies for all CAFE assays were reared on standard SY medium up until 22-24 hours before the CAFE assays, when they were anaesthetised with carbon dioxide (CO_2_) and sorted into groups of 5-10 and wet starved on 2% agar in water. For all assays, we chose to group-house flies in vials to more closely simulate their natural environments where they live in groups and to reduce noise of the data.

First, to test the effect of sex on food intake behaviour, male and female flies were reared to 7 days of age before performing assays. To understand the effect of mating status on food intake behaviour, newly eclosed flies were anaesthetized with CO_2_ and groups of 5 virgin females were transferred into narrow *Drosophila* vials (Genesee Scientific #:32-109) containing SY media for 5 days. The remaining males from the same cohort were maintained on SY media throughout this time. After 5 days, the virgin female flies were confirmed to be virgins from a lack of larvae in the vials. 5 males were then added to half of the total vials of virgin females and the flies were left to mate for 3 days. After the 3 days, flies were anaesthetised using CO_2_. The males were discarded, and the females were sorted into groups of either 5 mated or 5 virgin females. Finally, to investigate the effect of age, egg lays and collections were staged so that in 6 weeks’ time, there would be three cohorts of flies aged either 1-, 2-, 3-or 6-weeks old on the same date.

First, we investigated the effect of sex on food intake by comparing the food intake of 8-day-old males and females constrained to one of 3 P:C diets at low and high calories. We used the low-calorie diets for further experiments as the consistency of the high-calorie diets caused the food to accumulate in the capillary tubes. Next, to elucidate any effects of mating status, we compared the food intake of mated and virgin female flies. Finally, to reveal any effects of age, these assays were performed with groups of flies aged 1-, 2-, 3- and 6-weeks old.

For assays investigating the effect of sex on feeding behaviour, the number of replicates was 16-22 for both sexes and all diets, using vials with 6-10 flies. For assays using virgin and mated females, the number of replicates was 14-22 for both mating statuses and all diets, using vials of 3-10 flies. Finally, the number of replicates for assays on flies of different ages was 12-42 for each diet and all age groups, using 3-10 flies per vial.

All experiments were performed at 25°C and 60-80% humidity in a dark, controlled temperature room. 5 µL calibrated microcapillary tubes (Miles Scientific) were filled with one of three P:C diets (**Table 1**). A permanent marker was used to draw a line on the tube, indicating the initial total volume of food. The microcapillary tubes were then wrapped in cotton wool and plugged into the top of narrow vials (Genesee Scientific; Product Number: 32-116) containing 3-10 flies on 2% agar in water. Flies were then allowed to feed for 2 hours. Preliminary CAFE assays at 3 hours resulted in complete depletion of food in the capillary tube for many of the groups of mated females. Hence, we truncated these feeding sessions to 2 hours. To control for evaporation, capillary tubes were filled with one of the three diets and placed into vials that did not contain any flies. The amount of food that had evaporated was also recorded after 2 hours (**Supplementary Figure 1**).

### Quantification of food intake

A standard ruler was used to measure the difference in volume of food in the capillary tube before and after the flies fed. We used calibrated capillary tubes so that we could convert the amount of food consumed from mm into µL. The amount of food eaten per fly was calculated by dividing the total food consumed by the number of flies in each vial. Any flies that had died during the assay or were unable to feed (e.g. wings or body became stuck in media) were discounted. The total protein consumed per vial was calculated by multiplying the total intake by the concentration of protein in the diet. Similarly, the total carbohydrate intake was calculated by multiplying the total intake from the vial by the concentration of carbohydrate in the diet. These amounts were then divided by the total number of flies in the vial to generate an average protein and average carbohydrate intake per fly.

### Statistical analysis

All statistical analyses were performed using RStudio (version 4.1.2, 2021-11-01). To understand the differences in food intake when constrained to each diet for male and female flies and flies of different ages, we log transformed both P:C ratio and food intake, as diets weren’t equidistant and the data was not normally distributed. We fitted this using linear mixed effect models including diet, sex, and age as fixed effects, including all possible interactions, and date as a random block effect. To reveal significant effects of mating status on food intake, we again fitted the data using linear mixed effect models, with diet and mating status as fixed effects and including all possible interactions, and with date as a random block effect.

To test for differences in variance in protein and carbohydrate intake, we transformed our data into a set of normalised differences from the median of each data point, similar to the approach outlined in Carvalho and Mirth (2017). To do this, we calculated /*(x*_*ij*_ *- x*_*i*_.)/, where *x*_*ii*_ is the measured variable from the *j*th case from the *i*th group and *x*_*i*_ is the median for the *i*th group for each macronutrient. Data was fit with a generalised linear model, assuming a quasipoisson distribution. We then used estimated marginal means (from the emmeans package (Length, 2020)) to test for differences in the variance in macronutrient intake across macronutrient types and between flies of different sex, mating status, and age. Statistical significance was taken at *p* < 0.05.

### Ethical note

Flies were reared and housed at controlled densities in 300 ml bottles at 25°C and 80% relative humidity. All flies were provided with ∼70 ml appetitive food *ad libitum* at the bottom of their bottles. Flies were housed in populations of ∼100-200 flies per bottle and were not overcrowded. Cardboard bridges were placed in the bottles for enrichment and to reduce humidity - so that the flies remained dry and their ability to fly was not impeded. Flies were never starved for longer than 24 hours and were provided with a source of water during starvation (2% agar in water at the base of vial) to prevent effects of water stress (Fanson et al., 2012). Flies were lightly anaesthetised with CO_2_ when handling and were manipulated gently with a fine paintbrush. To cull flies, they were anaesthetised with CO_2_ before being tipped into 80% ethanol.

## RESULTS

### The effect of sex on food intake across P:C ratios

CAFE assays revealed a negative relationship between the P:C ratio of the diet and the food intake of both male and female 8-day-old flies (**Figure 1a; Table 2**), with flies consuming more food on the lowest P:C diet and less food on the highest P:C diet (**Figure 1a; Table 2**). Further, females had significantly higher overall food intake than males across all diets (**Figure 1a; Table 2**). Both male and female flies consumed significantly more food on the low-calorie diet than when on the high calorie diet (**Figure 1a; Table 2**).

**Figure 1:**
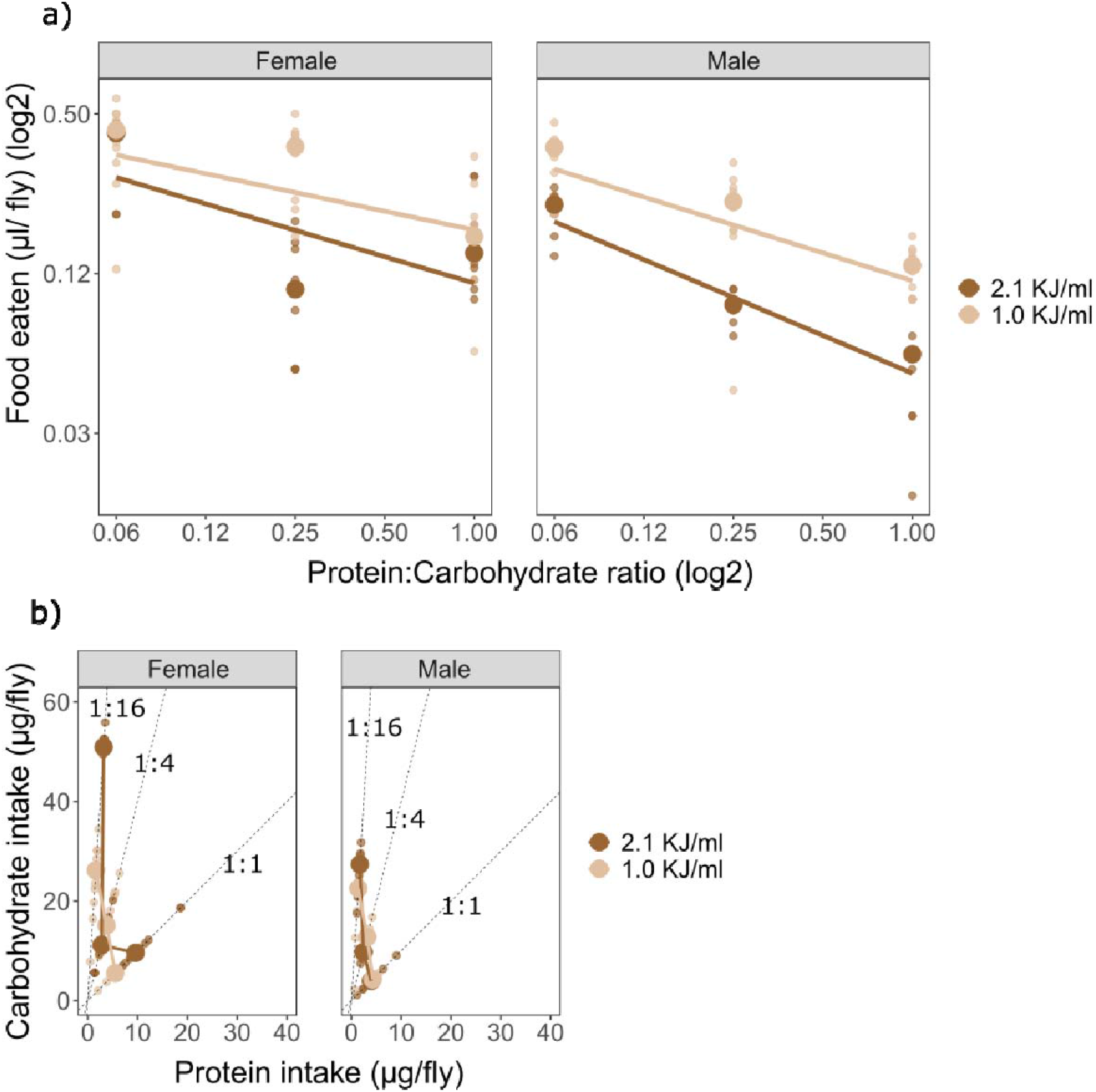
Male and female food intake across 3 P:C diets. A) Male and female flies demonstrated a negative relationship between P:C ratio and food intake. This pattern persisted across both high (2.1 KJ/ml) and low-calorie (1.0 KJ/ml) diets, but with flies on lower energy diets consuming more food than those on higher energy diets. B) Plotting the average carbohydrate intake per fly against the average protein intake per fly revealed that both male and female flies varied their carbohydrate intake more than their protein intake. Solid lines connect median protein/carbohydrate intake across ratios. Dashed lines represent the P:C ratios of the three diets (1:1 P:C, 1:4 P:C, and 1:16 P:C). Large circles = medians; small circles = replicates (average value of a group of 6 - 10 flies).

**Table 2.** ANOVA results for the effects of P:C ratio, sex and calories on the food intake of 8-day-old flies.

| Predictor | X<sup>2</sup> | df | p value | Significance |
| --- | --- | --- | --- | --- |
| P:C Ratio | 119.60 | 1 | <2.2e<sup>-16</sup> | *** |
| Calories | 20.33 | 1 | 6.53e<sup>-06</sup> | *** |
| Sex | 30.35 | 1 | 3.60e<sup>-08</sup> | *** |
| P:C Ratio*Calories | 0.65 | 1 | 0.42 | ns |
| P:C Ratio*Sex | 0.08 | 1 | 0.77 | ns |
| Calories*Sex | 0.89 | 1 | 0.35 | ns |
| P:C<br>Ratio*Calories*Sex | 0.73 | 1 | 0.39 | ns |

\*\*\* < 0.001; \*\* < 0.01; \* < 0.05; ns = not significant

To test for differences in variation in macronutrient intake between sexes, we compared the variation in carbohydrate and protein intake between male and female flies using our data set of normalised differences from the median of each data point. We found that both sexes varied their carbohydrate intake significantly more than their protein intake, and that female flies varied their carbohydrate intake more than males (**Figure 1b; Table 3**). However, the two sexes did not differ in their variation of protein intake. Given that the energy density of both nutrients is equal, this means that flies tend to more tightly control protein intake than carbohydrate intake.

**Table 3.**
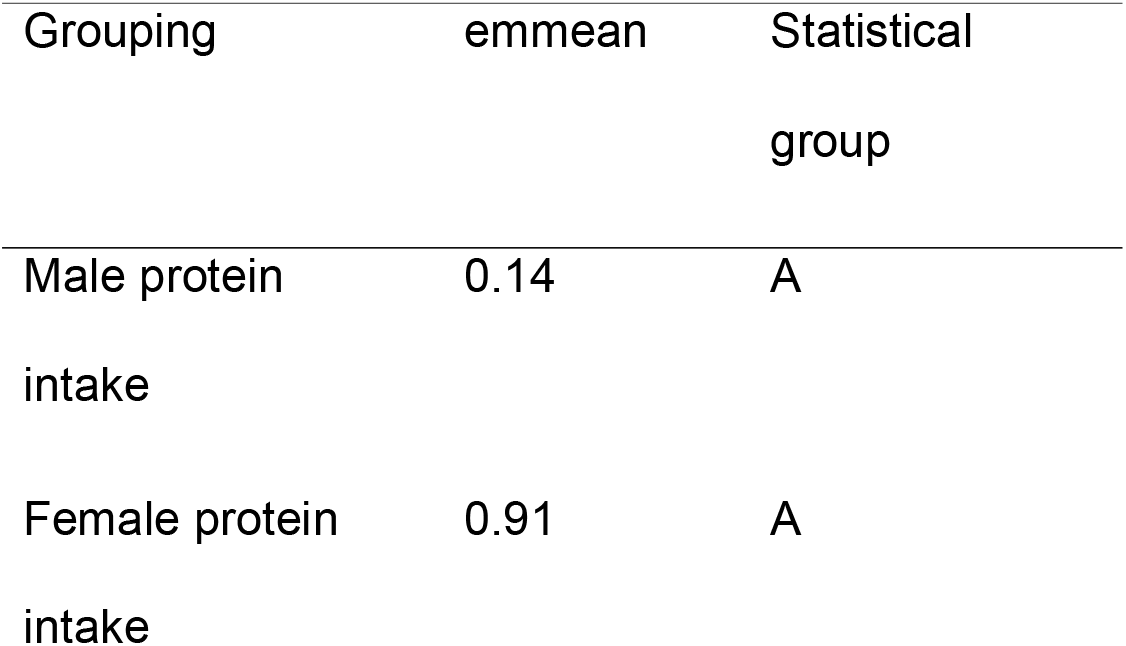
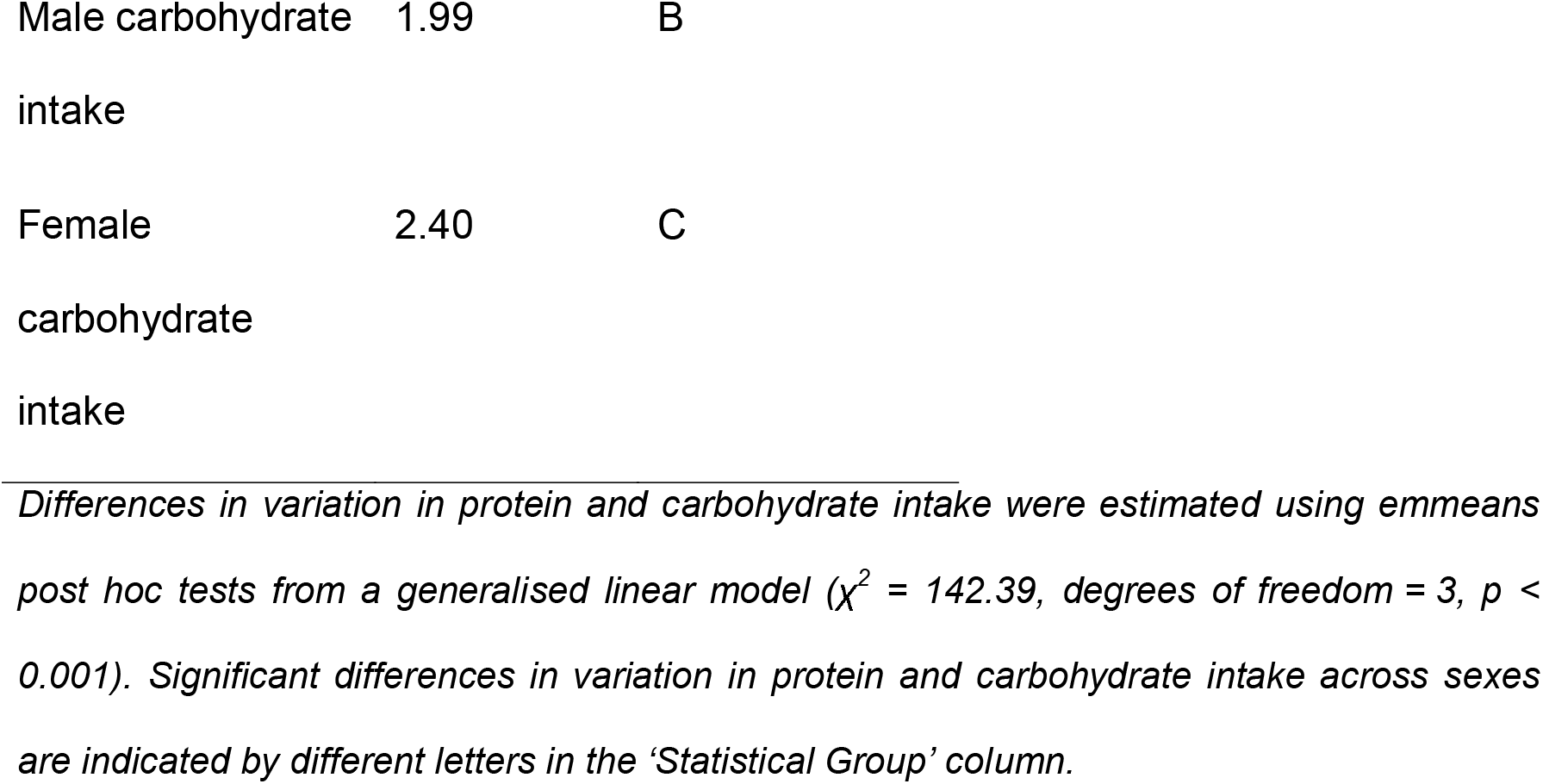
Both males and females show higher dispersion for carbohydrate intake than for protein intake.

| Grouping | emmean | Statistical<br>group |
| --- | --- | --- |
| Male protein<br>intake | 0.14 | A |
| Female protein<br>intake | 0.91 | A |
| Male carbohydrate intake | 1.99 | B |
| Female carbohydrate intake | 2.40 | C |

Differences in variation in protein and carbohydrate intake were estimated using emmeans post hoc tests from a generalised linear model ( $\chi^2 = 142.39$ , degrees of freedom = 3, $p < 0.001$ ). Significant differences in variation in protein and carbohydrate intake across sexes are indicated by different letters in the ‘Statistical Group’ column.

### The effect of mating status on food intake across P:C ratios

To uncover effects of mating status on food intake behaviour, we performed CAFE assays comparing 8-day-old virgin and mated females constrained to one of three diets. We found mating status significantly affected food intake, with mated female flies consuming significantly more food than virgin female flies (**Figure 2a**; **Table 4**). Further, post hoc comparisons revealed that both virgin and mated females varied their carbohydrate consumption significantly more than their protein consumption (**Figure 2b**; **Table 5**). Hence, while mated females ate more overall, virgin and mated female flies compromised for protein deficit in a similar manner.

**Figure 2:**
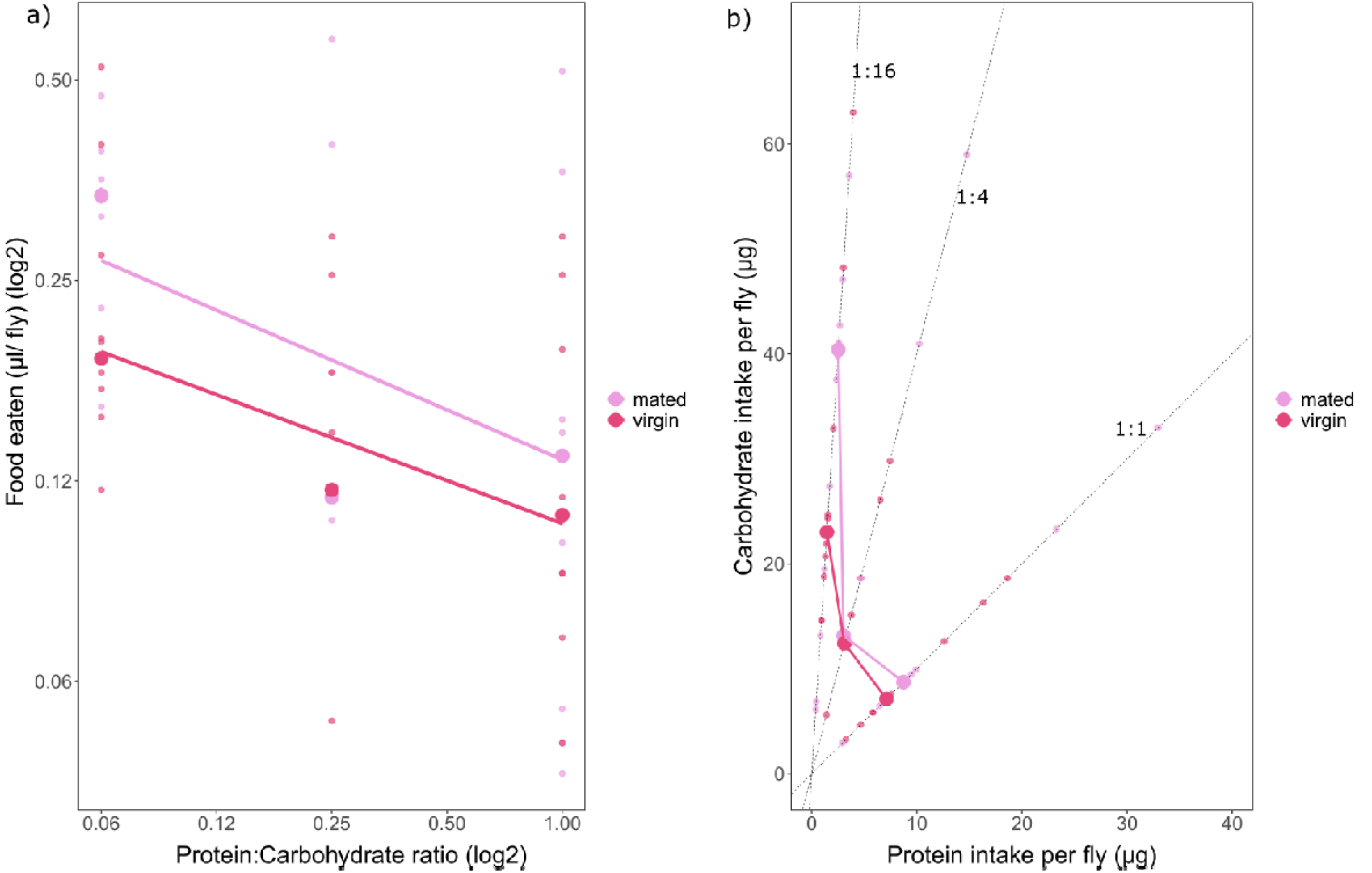
Virgin and mated 8-day-old female food intake across 3 P:C diets. A) Both virgin and mated female flies compromise for a protein deficit by consuming more food on the low P:C diet, but mated females consume more overall food (*p* <0.01). B) Carbohydrate and protein intake of virgin and mated females. Flies varied their carbohydrate intake significantly more than their protein intake regardless of mating status. Solid lines connect median protein/carbohydrate intake across ratios. Dashed lines represent diets (1:1 P:C, 1:4 P:C, and 1:16 P:C). Large circles = medians; small circles = replicates (average value of a group of 3-10 flies).

**Table 4.**
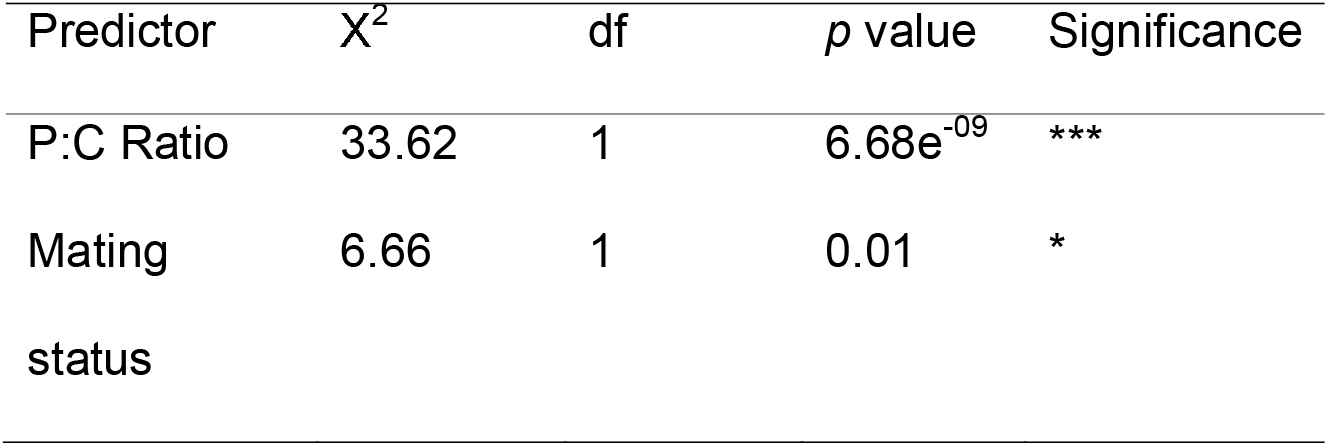
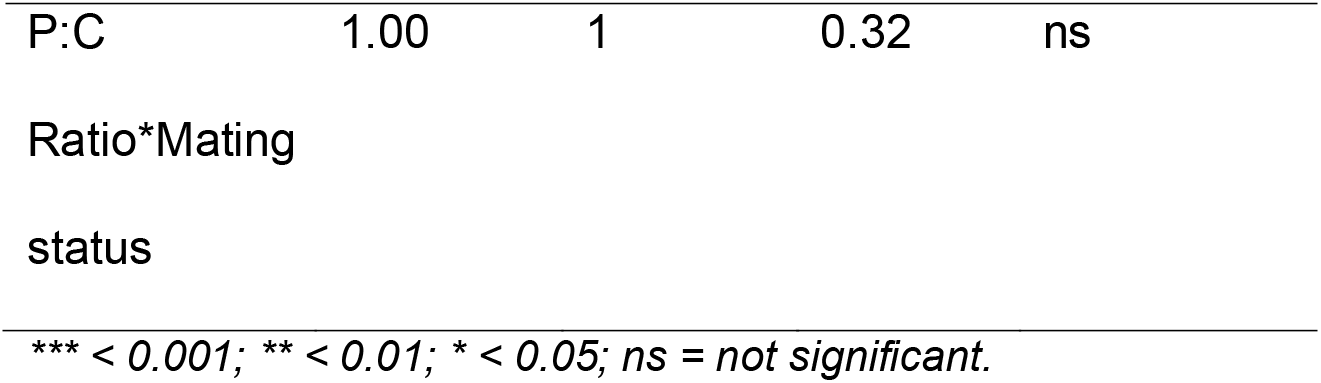
ANOVA results for the effects of P:C ratio and mating status on the food intake of 8-day-old flies.

**Table 5.**
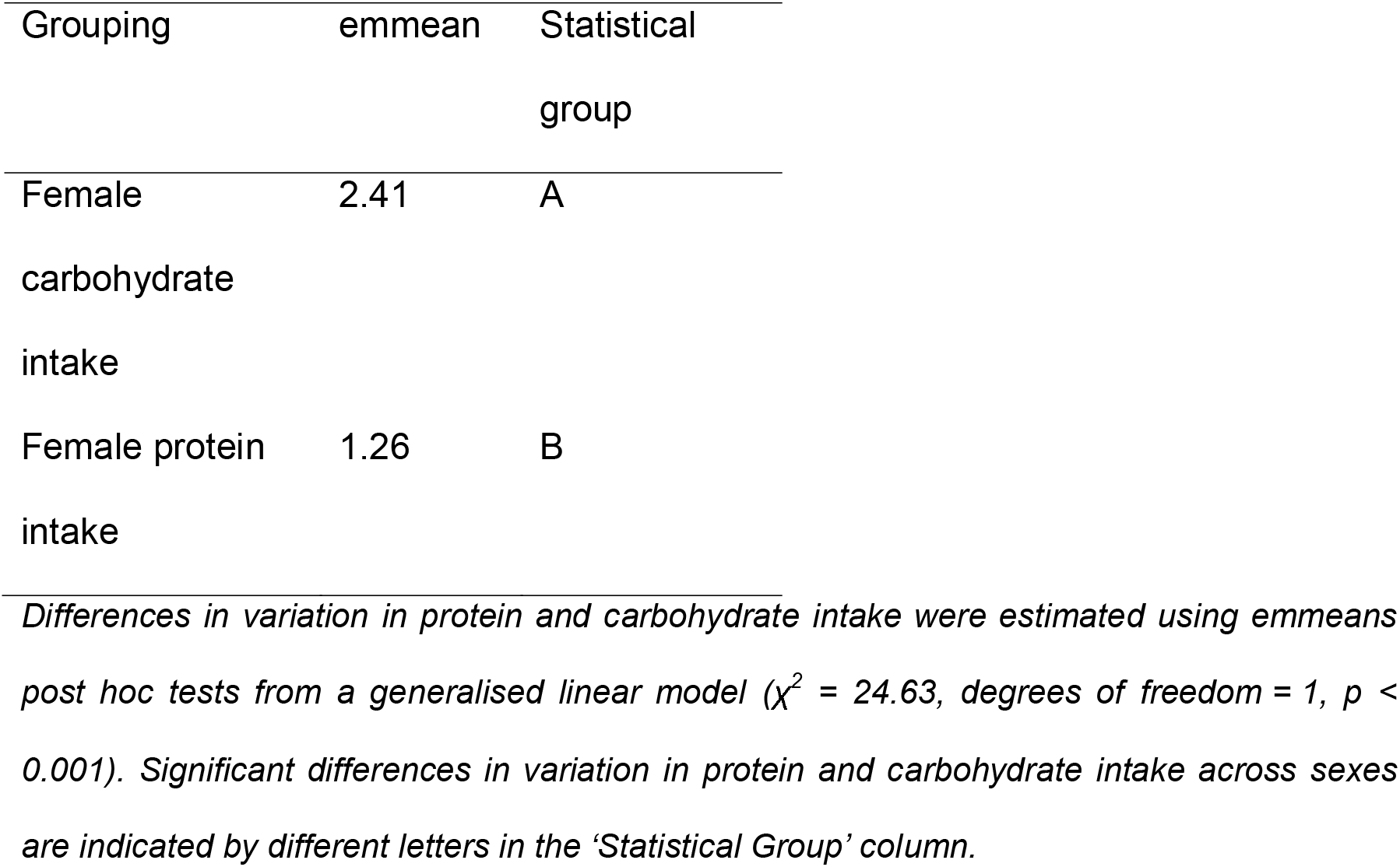
Females (mated and virgin combined) show higher dispersion in carbohydrate intake than in protein intake.

### The effect of age on food intake across P:C ratios

We next tested whether the capacity to compromise for protein deficit changed with age. To do this, we performed CAFE assays with flies at 1-, 2-, 3- and 6-weeks of age constrained to one of three diets. Consistent with previous assays, 1-week old flies demonstrated an ability to compromise for a protein deficit by consuming more food on diets with a low P:C ratio and less food on diets with a high P:C ratio. However, as flies aged, we found that they ate progressively less food overall (**Figure 3**). Further, while food intake decreased with increasing P:C ratio in 1-week old flies, we found this relationship changed with age (**Figure 3a**). We found significant interaction terms between P:C ratio and age, and sex and age (**Table 6**). Also, we uncovered a significant three-way interaction term between P:C ratio, age, and sex (**Table 6)**. This revealed that as flies age, the manner in which they adjust their food intake with the P:C ratio of their diet changes, and that the rate of change differs across sexes. Specifically, females from two weeks of age consume less overall food than their 1-week-old counterparts, and do not change their carbohydrate intake with the P:C ratio of the diet at this age, suggesting females do not compromise for a protein deficit from 2 weeks of age (**Figure 3**; **Table 7-8**). For males, this transition occurs later – sometime after 3 weeks of age; with the greatest food consumption on low P:C diets at two weeks of age and significantly higher consumption of carbohydrates compared to protein until sometime after 3-weeks of age (**Figure 3**; **Table 7; Table 9**).

**Figure 3:**
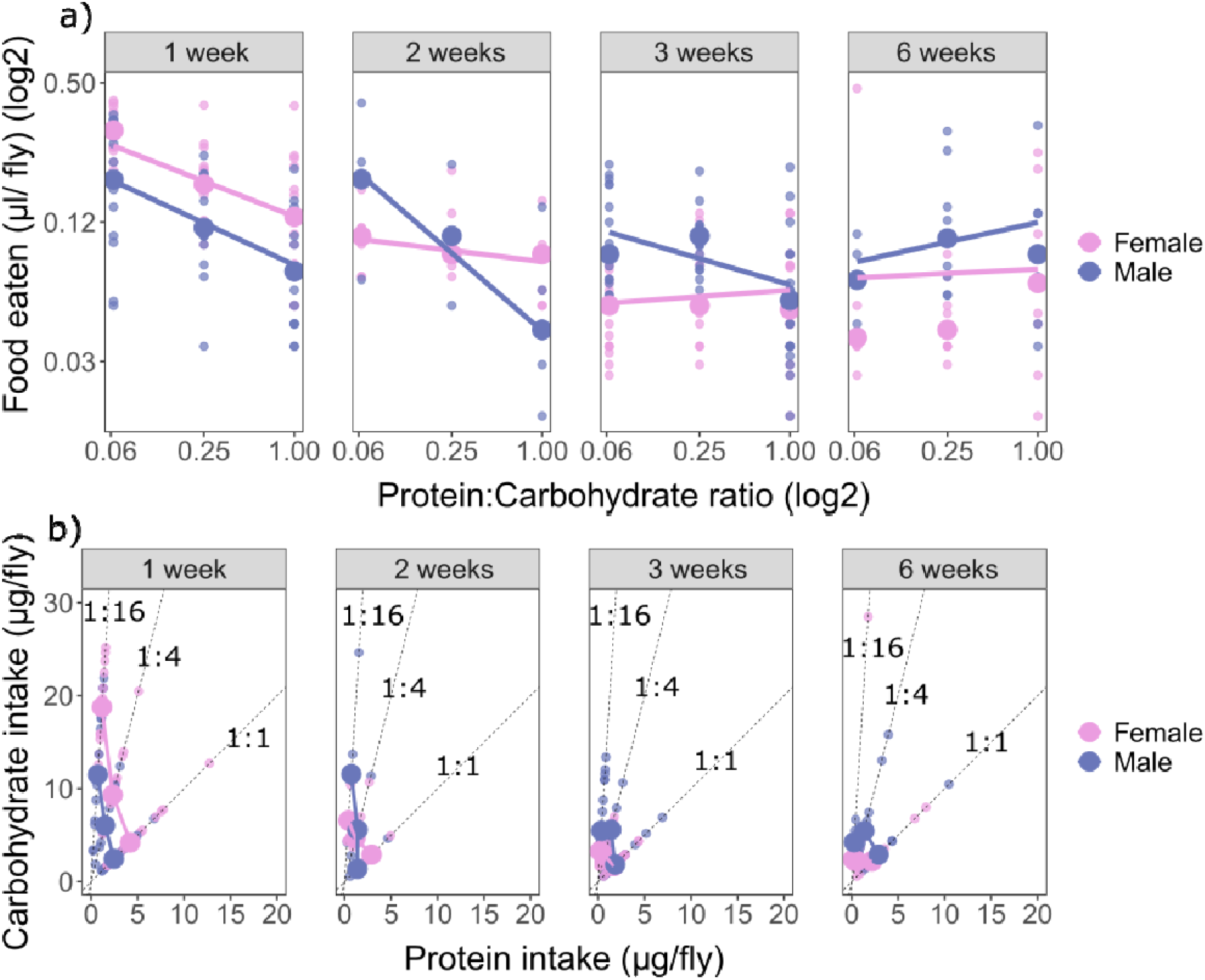
Food intake and the rules of compromise change with age in both males and females. A) Both male and female flies compromise for a protein deficit by consuming more food on the low P:C diet at 1-week of age. However, from 2-weeks onwards, female flies cease to alter their intake across diets. Male flies continue to alter their intake across diets until at least 3 weeks of age. B) 1-week-old mated females consume more protein and carbohydrates than males. Solid lines connect median protein/carbohydrate intake across ratios. Dashed lines represent diets (1:1 P:C, 1:4 P:C, and 1:16 P:C). Large circles = medians; small circles = replicates (average value of a group of 3-10 flies).

**Table 6.**
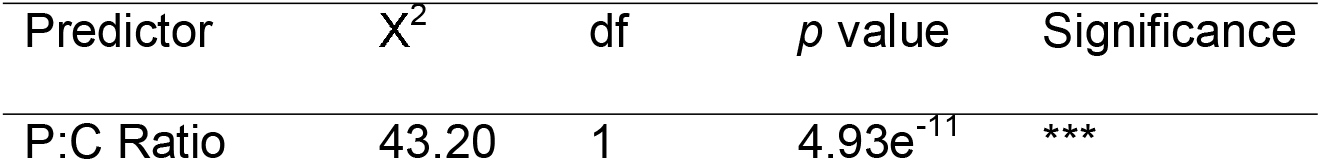
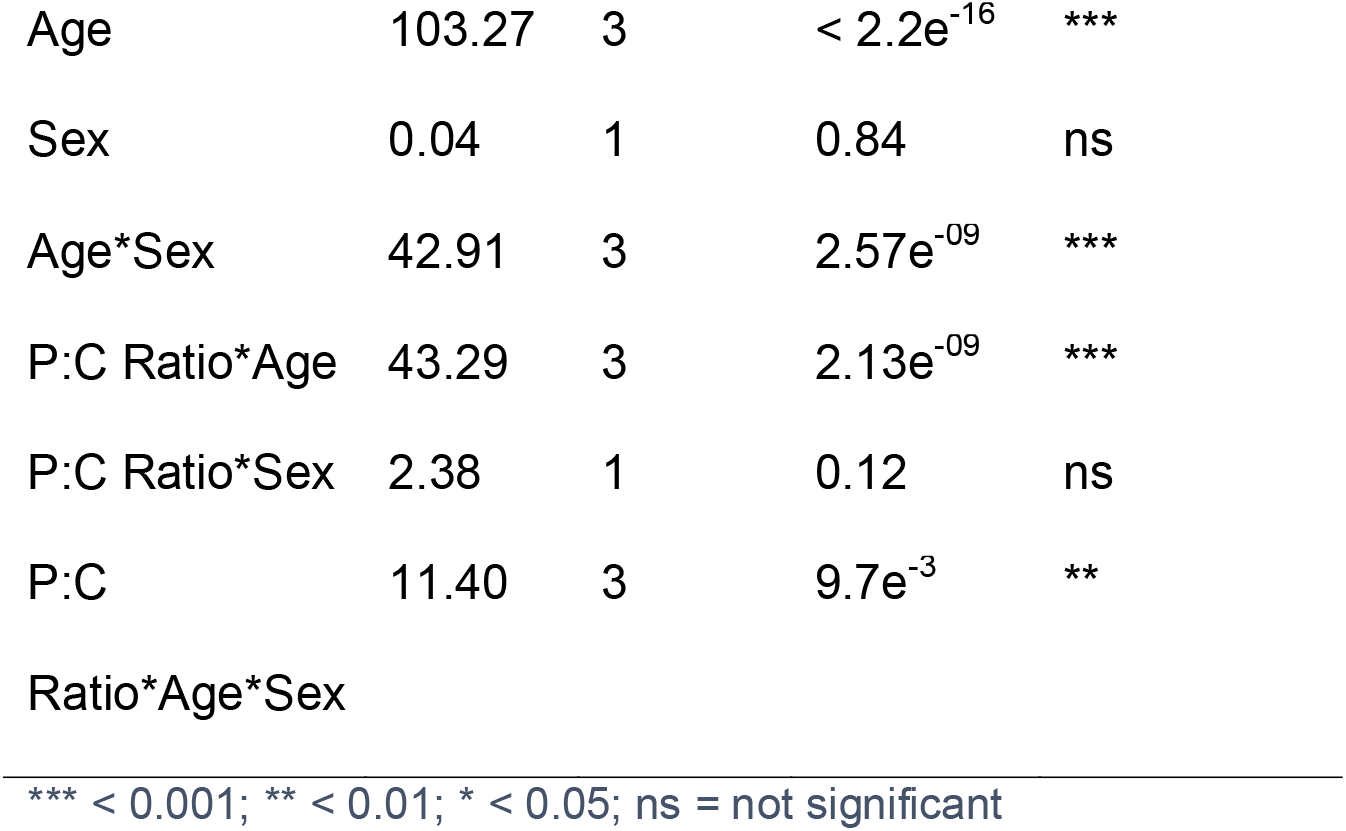
ANOVA results for the effects of P:C ratio, age and sex on the food intake of 1-, 2-, 3- and 6-week-old flies.

**Table 7.** Differences in overall food intake in 1-, 2, 3-, and 6-week-old mated flies.

|  |  | emtrend | Statistical<br>group |
| --- | --- | --- | --- |
| Female | 1-week-old | -0.10 | A |
|  | 2-week-old | -0.02 | B |
|  | 3-week-old | 0.01 | B |
|  | 6-week-old | 0.01 | B |
| Male | 1-week-old | -0.08 | A |
|  | 2-week-old | -0.10 | B |
|  | 3-week-old | -0.03 | C |
|  | 6-week-old | 0.03 | D |

Differences in food intake with age in females and males were estimated using emmeans post hoc tests from a generalised linear model in Table 7. Significant differences in food intake between weeks are indicated by different letters in the 'Statistical Group' column.

**Table 8.** Differences in the dispersion in protein and carbohydrate intake between 1-, 2, 3-, and 6-week-old mated female flies.

|  |  | Grouping | emmean | Statistical group |
| --- | --- | --- | --- | --- |
| Female | 1-week-old | Protein intake | 0.53 | A |
|  |  | Carb intake | 2.07 | B |
|  | 2-week-old | Protein intake | -0.76 | A |
|  |  | Carb intake | 0.71 | A |
|  | 3-week-old | Protein intake | -0.21 | A |
|  |  | Carb intake | 0.40 | B |
|  | 6-week-old | Protein intake | 0.25 | A |
|  |  | Carb intake | 0.95 | B |

Differences in variation in protein and carbohydrate intake were estimated using emmeans post hoc tests from a generalised linear model ( $\chi^2 = 126.83$ , degrees of freedom = 3, $p < 0.001$ ). Significant differences in variation in protein and carbohydrate intake across sexes are indicated by different letters in the 'Statistical Group' column.

**Table 9.** Differences in the dispersion in protein and carbohydrate intake between 1-, 2, 3-, and 6-week-old male flies.

|  |  | Grouping | emmean | Statistical group |
| --- | --- | --- | --- | --- |
| Male | 1-week-old | Protein intake | -0.04 | A |
|  |  | Carb intake | 1.50 | B |
|  | 2-week-old | Protein intake | -0.51 | A |
|  |  | Carb intake | 1.42 | B |
|  | 3-week-old | Protein intake | -0.31 | A |
|  |  | Carb intake | 0.85 | B |
|  | 6-week-old | Protein intake | 0.32 | A |
|  |  | Carb intake | 0.88 | A |

Differences in variation in protein and carbohydrate intake were estimated using emmeans post hoc tests from a generalised linear model ( $\chi^2 = 126.83$ , degrees of freedom = 3, $p < 0.001$ ). Significant differences in variation in protein and carbohydrate intake across sexes are indicated by different letters in the 'Statistical Group' column.

Combined with the food intake analysis above, this suggests that while older flies still appear to adjust their food intake with the P:C ratio of the food, the degree of protein leveraging weakens as flies age. Further, females cease protein leveraging at a younger age than males.

## DISCUSSION

Altering the ratio of macronutrients in an animal’s diet has profound effects on its reproductive success, lifespan, and overall health (Simpson & Raubenheimer, 2012). Consequently, animals must regulate their food intake to reach nutritional targets necessary to sustain fitness. Since natural selection exerts greater pressure early in life (Jiang & Assis, 2017; Rose et al., 2007), regulating nutrient intake should be most important during growth and early reproduction. In later life, there are signs that regulation of nutrient intake to sustain health is lost since malnutrition is particularly prevalent in older age groups (Shahar et al., 2002). Despite this, no studies to date have investigated changes in nutrient intake across age groups in *Drosophila*. In this study, we used CAFE assays to discern how sex, mating status, and age affect the way in which *D. melanogaster* regulates its nutrient intake. We found a sex-specific, age-dependent change in macronutrient intake. This study afforded novel understanding of how food intake behaviours change with age, providing a springboard for future research into the potential role of the regulation of macronutrient intake in the anorexia of ageing.

*D. melanogaster* larvae consume more food on low P:C diets and less food on high P:C diets (Carvalho & Mirth, 2017). We hypothesized that adult fruit flies would regulate their food intake in a different manner, since their life stage requires nutrients tailored to support reproduction and dispersal, while the larval stage requires nutrients for growth and development (Nestel et al., 2016). However, we found that both young male and young female adult flies consumed more food on the low P:C diet, compromising for the protein deficit and exhibiting protein leveraging behaviour, akin to their larval counterparts. Hence, both as larvae and as young adults *D. melanogaster* consumed more food on a low P:C diet, consistent with flies consuming more food to reach a protein target. Fanson et al. (2012) found that young adult *Drosophila* prioritize intake of both carbohydrates and protein equally when provided with a choice between water and diets with varying yeast and sucrose content. However, plotting the protein and carbohydrate intake per fly in the current study showed that young adult flies more tightly regulated their protein intake, while carbohydrate intake varied more widely. This was further confirmed statistically by post hoc comparisons which revealed that both young male and female flies kept their protein intake constant across diets while significantly varying their carbohydrate intake. This finding provides yet more evidence that, like mice and humans, flies also use protein leveraging rules of compromise when their diet does not allow them to reach their intake target (Simpson & Raubenheimer, 2005).

The overconsumption of energy due to protein leveraging is a compelling explanation for some instances of obesity in children, adolescents (Saner et al., 2020), and adults (Simpson & Raubenheimer, 2005). Many of the highly processed foods that are abundant in society today are diluted in protein and, as a result, have a higher percentage of fat and carbohydrates. Individuals will consume more total energy from these highly processed foods to compensate for a diet diluted in protein, and because of the highly appetitive nature of the dilutants (Steele et al., 2018). Our data provides additional evidence that the protein leveraging phenomena is prevalent across taxa.

Virgin and mated females both balanced their nutrient intake similarly. It has been previously posited that flies’ feeding behaviour can anticipate future states (Walker et al., 2017). Our data offers evidence for anticipatory regulation of feeding behaviour since the virgin females behaved as if they were mated, in terms of macronutrient balancing. Consistent with previous research, we found that young mated females consumed more overall food than young virgin females (Camus et al., 2018). This increased food consumption in post-copulatory females has been attributed to an increased protein demand for egg production and maintenance in *Drosophila* species (Bellutti et al., 2018; Good & Tatar, 2001; Simmons & Bradley, 1997; Terashima & Bownes, 2004).

We demonstrated a sex-specific, age-dependent effect on food intake behaviour in *D. melanogaster*. As flies aged, they ceased to adjust their feeding rates with the P:C content of the diet. We found that female flies showed weaker protein leveraging behaviour at 2 weeks of age, while protein leveraging declined more slowly in males. When given the option to self-compose a diet, male flies choose a diet containing four times more carbohydrate than protein (Lee et al., 2013). This is likely attributed to male reproductive success relying upon courtship activities that are energetically demanding (Partridge, Ewing, et al., 1987). However, in the current study males compensated for protein dilution for longer than females. Protein in the diet is crucial for both male and female *Drosophila* reproductive success (Fricke et al., 2008). Our findings suggest that male flies utilise protein for reproductive success across their entire lifetime, and not only in their youth.

When constrained to a low P:C diet in the present study, female mated *D. melanogaster* overconsumed carbohydrates and overall food to compromise for the protein deficit and to reach their protein intake target. However, as the flies aged, they no longer consumed more food on the low protein diet. Previous research has shown that when given a choice between diets, young mated female flies self-select a diet high in protein (Lee et al., 2013). Diets high in protein relative to carbohydrates maximise egg-laying rate in young females, while diets low in protein relative to carbohydrates extend lifespan (Lee et al., 2008). Our data demonstrates that as female flies age, they no longer prioritise reaching their protein intake target for egg-laying via protein leveraging. Hence, female *D. melanogaster* appear to prioritise egg-laying over lifespan only at a young age.

Increased protein consumption results in increased egg laying in female flies (Kim et al., 2020; Lee, 2015; Lee et al., 2008; Partridge, Green, et al., 1987). Our finding that females consumed the most protein while young is consistent with data showing that female flies lay the greatest number of eggs in their first 10 days of life (Good & Tatar, 2001). Our results for the effects of mating status and age on food intake behaviour suggest that mated females consume an abundance of nutrients at a young age that provides them with resources for increased egg production while they are young. Then, as the females age, they decrease egg production and subsequently their drive to eat protein diminishes.

Olfaction and feeding behaviour are intertwined. Odours provide the location of food to foraging animals and can alter the responses of hunger-sensing neurons (Chen et al., 2015). Further, removing sense of smell has been shown to protect against diet-induced obesity in rodents (Riera et al., 2017). As flies age, they experience functional declines in olfaction and locomotion (Cook-Wiens & Grotewiel, 2002). For instance, male *Bactrocera tyroni* (Queensland Fruit Fly) demonstrate a significant drop in olfactory response at 15 weeks of age; females exhibit this earlier at 6 weeks of age (Tasnin et al., 2020). Additionally, a decline in olfactory performance has been reported in *Drosophila* in their first few weeks of life, with female olfactory performance declining more rapidly (Simon et al., 2006). In fact, as humans age, they also experience a decline in olfactory function which leads to appetite loss, decreased enjoyment of food, more food dislikes, and an altered diet (Wysocki & Pelchat, 1993). It is plausible in our study that the flies’ diminishing olfactory ability resulted in poor nutrient regulation as they aged, and that the earlier decline in protein leveraging behaviour in females mirrors their earlier loss of olfactory function. Hence, olfaction may play a necessary function in the regulation of nutrient intake.

One caveat of this study is that yeast contains a variety of nutrients. Hence, altering the concentration of yeast across diets may have resulted in changes in the quantity of micronutrients in the diet, namely vitamins and minerals. For stricter control of micronutrients, future studies may endeavour to use synthetic diets such as those designed by Piper et al. (2014). A second caveat worth discussing is that the low P:C diet may have been more palatable than the high P:C diets, as it contained more sucrose and tasted sweeter. Hence, performing the CAFE assays for longer durations may disentangle any phagostimulant effects of the diet from actual homeostatic regulation of protein intake. Finally, since this study did not allow the flies to self-compose a diet, we were unable to calculate intake targets for our flies specifically. As such, an alternative explanation for the change in nutrient intake with age could be that the flies’ intake targets change with age.

From our study, we have provided new findings about how *Drosophila* use rules of compromise when unable to reach their nutrient targets. Firstly, we have shown that adult *D. melanogaster* alter their food intake in the same manner as larvae - namely that both adults and larvae employ protein leveraging to compromise for low protein diets. Secondly, we have added to previous evidence that mated females consume more overall food than virgin females and mated males, however they appear to use the same rule of compromise. Finally, we found that protein leveraging declines with age and diminishes faster in female flies than in males. This knowledge allows for future studies to probe into the neural mechanisms of *Drosophila* macronutrient balancing and to investigate the effects of external factors such as the flies’ surrounding environment and chemosensory ability. Inhibiting the olfactory system in flies across diets and with age would be invaluable in determining if a decline in olfactory function plays a role in the decline of macronutrient balancing that we have observed in the current study.

## AUTHOR CONTRIBUTIONS

**Helen J Rushby**: Conceptualization, data curation, formal analysis, investigation, methodology, validation, resources, writing – original draft, visualization, project administration. **Christen K Mirth**: Conceptualization, methodology, formal analysis, data curation, writing – review and editing, supervision, project administration, funding acquisition. **Matthew D W Piper**: Conceptualization, methodology, writing – review and editing, supervision, project administration, funding acquisition. **Zane B Andrews**: Conceptualization, writing – review and editing, supervision, funding acquisition.

## ACKNOWLEDGEMENTS

We would like to thanks the members of the Mirth and Piper lab, past and present, for their help in this project. In particular, we would like to thank Dr Jade Kannangara for maintaining the flies during COVID-19 lockdowns. We would also like to thank Dr Avishikta Chakraborty for assisting with data analysis.

## DATA AVAILABILITY

All data and scripts are available in online repositories on Figshare (DOI: 10.26180/19179113).

## REFERENCES

Bellutti, N., Gallmetzer, A., Innerebner, G., Schmidt, S., Zelger, R., & Koschier, E. H. (2018). Dietary yeast affects preference and performance in Drosophila suzukii. Journal of pest science, 91(2), 651–660.

Bowman, E., & Tatar, M. (2016). Reproduction regulates Drosophila nutrient intake through independent effects of egg production and sex peptide: implications for aging. Nutrition and healthy aging, 4(1), 55–61.

Camus, M. F., Huang, C.-C., Reuter, M., & Fowler, K. (2018). Dietary choices are influenced by genotype, mating status, and sex in Drosophila melanogaster. Ecology and Evolution, 8(11), 5385–5393. https://doi.org/10.1002/ece3.4055

Carvalho, M. J., & Mirth, C. K. (2017). Food intake and food choice are altered by the developmental transition at critical weight in Drosophila melanogaster. Animal Behaviour, 126, 195–208. https://doi.org/https://doi.org/10.1016/j.anbehav.2017.02.005

Chen, C., Lin, Y., Kuo, T., & Knight, K. (2015). Sensory detection of food rapidly modulates arcuate feeding circuits. Cell, 160, 829–841.

Cook-Wiens, E., & Grotewiel, M. S. (2002). Dissociation between functional senescence and oxidative stress resistance in Drosophila. Experimental Gerontology, 37(12), 1347–1357. https://doi.org/https://doi.org/10.1016/S0531-5565(02)00096-7

Fanson, B. G., Yap, S., & Taylor, P. W. (2012). Geometry of compensatory feeding and water consumption in Drosophila melanogaster. Journal of Experimental Biology, 215(5), 766–773. https://doi.org/10.1242/jeb.066860

Felton, A. M., Felton, A., Raubenheimer, D., Simpson, S. J., Foley, W. J., Wood, J. T., Wallis, I. R., & Lindenmayer, D. B. (2009). Protein content of diets dictates the daily energy intake of a freeranging primate. Behavioral Ecology, 20(4), 685–690.

Floegel, A., & Pischon, T. (2012). Low carbohydrate-high protein diets. British Medical Journal Publishing Group, 344, e3081. https://doi.org/https://doi.org/10.1136/bmj.e3801

Fricke, C., Bretman, A., & Chapman, T. (2008). Adult male nutrition and reproductive success in Drosophila Melanogaster. Evolution, 62(12), 3170–3177. https://doi.org/https://doi.org/10.1111/j.1558-5646.2008.00515.x

Good, T., & Tatar, M. (2001). Age-specific mortality and reproduction respond to adult dietary restriction in Drosophila melanogaster. Journal of Insect Physiology, 47(12), 1467–1473.

Gosby, A. K., Conigrave, A. D., Lau, N. S., Iglesias, M. A., Hall, R. M., Jebb, S. A., Brand-Miller, J., Caterson, I. D., Raubenheimer, D., & Simpson, S. J. (2011). Testing protein leverage in lean humans: a randomised controlled experimental study. PloS one, 6(10), e25929.

Ja, W. W., Carvalho, G. B., Mak, E. M., de la Rosa, N. N., Fang, A. Y., Liong, J. C., Brummel, T., & Benzer, S. (2007). Prandiology of Drosophila and the CAFE assay. Proceedings of the National Academy of Sciences, 104(20), 8253–8256. https://doi.org/10.1073/pnas.0702726104

Jensen, K., Mayntz, D., Toft, S., Clissold, F. J., Hunt, J., Raubenheimer, D., & Simpson, S. J. (2012). Optimal foraging for specific nutrients in predatory beetles. Proceedings of the Royal Society B: Biological Sciences, 279(1736), 2212–2218.

Jiang, X., & Assis, R. (2017). Natural Selection Drives Rapid Functional Evolution of Young Drosophila Duplicate Genes. Molecular Biology and Evolution, 34(12), 3089–3098. https://doi.org/10.1093/molbev/msx230

Kim, K., Jang, T., Min, K.-J., & Lee, K. P. (2020). Effects of dietary protein:carbohydrate balance on life-history traits in six laboratory strains of Drosophila melanogaster. Entomologia Experimentalis et Applicata, 168(6-7), 482–491. https://doi.org/https://doi.org/10.1111/eea.12855

Kremer, S., Mojet, J., & Kroeze, J. H. (2007). Differences in perception of sweet and savoury waffles between elderly and young subjects. Food Quality and Preference, 18(1), 106–116.

Lagiou, P., Sandin, S., Lof, M., Trichopoulos, D., Adami, H.-O., & Weiderpass, E. (2012). Low carbohydrate-high protein diet and incidence of cardiovascular diseases in Swedish women: prospective cohort study. The Bmj, 344, e4026.

Le Couteur, D. G., Solon-Biet, S., Cogger, V. C., Mitchell, S. J., Senior, A., de Cabo, R., Raubenheimer, D., & Simpson, S. J. (2016). The impact of low-protein high-carbohydrate diets on aging and lifespan. Cellular and Molecular Life Sciences, 73(6), 1237–1252. https://doi.org/10.1007/s00018-015-2120-y

Lee, K. P. (2015). Dietary protein:carbohydrate balance is a critical modulator of lifespan and reproduction in Drosophila melanogaster: A test using a chemically defined diet. Journal of Insect Physiology, 75, 12–19. https://doi.org/https://doi.org/10.1016/j.jinsphys.2015.02.007

Lee, K. P., Kim, J.-S., & Min, K.-J. (2013, length). Sexual dimorphism in nutrient intake and life span is mediated by mating in Drosophila melanogaster. Animal Behaviour, 86(5), 987–992. https://doi.org/https://doi.org/10.1016/j.anbehav.2013.08.018

Lee, K. P., Simpson, S. J., Clissold, F. J., Brooks, R., Ballard, J. W. O., Taylor, P. W., Soran, N., & Raubenheimer, D. (2008). Lifespan and reproduction in Drosophila: new insights from nutritional geometry. Proceedings of the National Academy of Sciences, 105(7), 2498–2503.

Length, R. V. (2020). emmeans: Estimated marginal means, aka least-squares means. R package version 152-1 https://CRANR-projectorg/package=emmeans.

Linford, N. J., Bilgir, C., Ro, J., & Pletcher, S. D. (2013). Measurement of lifespan in Drosophila melanogaster. JoVE (Journal of Visualized Experiments)(71), e50068.

Maklakov, A. A., Simpson, S. J., Zajitschek, F., Hall, M. D., Dessmann, J., Clissold, F., Raubenheimer, D., Bonduriansky, R., & Brooks, R. C. (2008). Sex-Specific Fitness Effects of Nutrient Intake on Reproduction and Lifespan. Current Biology, 18(14), 1062–1066. https://doi.org/https://doi.org/10.1016/j.cub.2008.06.059

Morley, J. E. (1997). Anorexia of aging: physiologic and pathologic. The American Journal of Clinical Nutrition, 66(4), 760–773. https://doi.org/10.1093/ajcn/66.4.760

Nestel, D., Papadopoulos, N. T., Pascacio-Villafán, C., Righini, N., Altuzar-Molina, A. R., & Aluja, M. (2016). Resource allocation and compensation during development in holometabolous insects. Journal of Insect Physiology, 95, 78–88.

Paddon-Jones, D., & Leidy, H. (2014). Dietary protein and muscle in older persons. Current opinion in clinical nutrition and metabolic care, 17(1), 5–11. https://doi.org/10.1097/MCO.0000000000000011

Partridge, L., Ewing, A., & Chandler, A. (1987). Male size and mating success in Drosophila melanogaster: the roles of male and female behaviour. Animal Behaviour, 35(2), 555–562. https://doi.org/https://doi.org/10.1016/S0003-3472(87)80281-6

Partridge, L., Green, A., & Fowler, K. (1987). Effects of egg-production and of exposure to males on female survival in Drosophila melanogaster. Journal of Insect Physiology, 33(10), 745–749. https://doi.org/https://doi.org/10.1016/0022-1910(87)90060-6

Piper, M. D., Blanc, E., Leitão-Gonçalves, R., Yang, M., He, X., Linford, N. J., Hoddinott, M. P., Hopfen, C., Soultoukis, G. A., & Niemeyer, C. (2014). A holidic medium for Drosophila melanogaster. Nature Methods, 11(1), 100.

Raubenheimer, D., & Simpson, S. J. (1997). Integrative models of nutrient balancing: application to insects and vertebrates. Nutrition research reviews, 10(1), 151–179.

Ribeiro, C., & Dickson, B. J. (2010). Sex Peptide Receptor and Neuronal TOR/S6K Signaling Modulate Nutrient Balancing in Drosophila. Current Biology, 20(11), 1000–1005. https://doi.org/https://doi.org/10.1016/j.cub.2010.03.061

Riera, C. E., Tsaousidou, E., Halloran, J., Follett, P., Hahn, O., Pereira, M. M. A., Ruud, L. E., Alber, J., Tharp, K., Anderson, C. M., Brönneke, H., Hampel, B., Filho, C. D. d. M., Stahl, A., Brüning, J. C., & Dillin, A. (2017). The Sense of Smell Impacts Metabolic Health and Obesity. Cell metabolism, 26(1), 198–211. https://doi.org/https://doi.org/10.1016/j.cmet.2017.06.015

Roberts, S. B. (2000, 2000). Regulation of energy intake in older adults: recent findings and implications. The journal of nutrition, health & aging, 4(3), 170–171.

Rodrigues, M. A., Martins, N. E., Balancé, L. F., Broom, L. N., Dias, A. J. S., Fernandes, A. S. D., Rodrigues, F., Sucena, É., & Mirth, C. K. (2015). Drosophila melanogaster larvae make nutritional choices that minimize developmental time. Journal of Insect Physiology, 81, 69–80. https://doi.org/https://doi.org/10.1016/j.jinsphys.2015.07.002

Rose, M. R., Rauser, C. L., Benford, G., Matos, M., & Mueller, L. D. (2007). Hamilton’s forces of natural selection after forty years. Evolution, 61(6), 1265–1276. https://doi.org/https://doi.org/10.1111/j.1558-5646.2007.00120.x

Saner, C., Tassoni, D., Harcourt, B. E., Kao, K. T., Alexander, E. J., McCallum, Z., Olds, T., Rowlands, A. V., Burgner, D. P., Simpson, S. J., Raubenheimer, D., Senior, A. M., Juonala, M., & Sabin, M. A. (2020). Evidence for Protein Leveraging in Children and Adolescents with Obesity. Obesity, 28, 822–829. https://doi.org/10.1002/oby.22755

Shahar, S., Chee, K. Y., & Wan Chik, W. C. P. (2002). Food intakes and preferences of hospitalised geriatric patients. BMC Geriatrics, 2(1), 3. https://doi.org/10.1186/1471-2318-2-3

Simmons, F. H., & Bradley, T. J. (1997). An analysis of resource allocation in response to dietary yeast in Drosophila melanogaster. Journal of Insect Physiology, 43(8), 779–788. https://doi.org/https://doi.org/10.1016/S0022-1910(97)00037-1

Simon, A. F., Liang, D. T., & Krantz, D. E. (2006). Differential decline in behavioral performance of Drosophila melanogaster with age. Mechanisms of Ageing and Development, 127(7), 647–651. https://doi.org/https://doi.org/10.1016/j.mad.2006.02.006

Simpson, S. J., Batley, R., & Raubenheimer, D. (2003). Geometric analysis of macronutrient intake in humans: the power of protein? Appetite, 41(2), 123–140.

Simpson, S. J., Le Couteur, D. G., & Raubenheimer, D. (2015). Putting the Balance Back in Diet. Cell, 161(1), 18–23. https://doi.org/https://doi.org/10.1016/j.cell.2015.02.033

Simpson, S. J., & Raubenheimer, D. (2005). Obesity: the protein leverage hypothesis. obesity reviews, 6(2), 133–142.

Simpson, S. J., & Raubenheimer, D. (2012). The Nature of Nutrition: A Unifying Framework from Animal Adaptation to Human Obesity. Princeton University Press. https://books.google.com.au/books?id=dzRFczYo8xQC

Simpson, S. J., Sibly, R. M., Lee, K. P., Behmer, S. T., & Raubenheimer, D. (2004). Optimal foraging when regulating intake of multiple nutrients. Animal Behaviour, 68(6), 1299–1311. https://doi.org/https://doi.org/10.1016/j.anbehav.2004.03.003

Solon-Biet, Samantha M., McMahon, Aisling C., Ballard, J. William O., Ruohonen, K., Wu, Lindsay E., Cogger, Victoria C., Warren, A., Huang, X., Pichaud, N., Melvin, Richard G., Gokarn, R., Khalil, M., Turner, N., Cooney, Gregory J., Sinclair, David A., Raubenheimer, D., Le Couteur, David G., & Simpson, Stephen J. (2014). The Ratio of Macronutrients, Not Caloric Intake, Dictates Cardiometabolic Health, Aging, and Longevity in Ad Libitum-Fed Mice. Cell metabolism, 19(3), 418–430. https://doi.org/https://doi.org/10.1016/j.cmet.2014.02.009

Sørensen, A., Mayntz, D., Raubenheimer, D., & Simpson, S. J. (2008). Protein-leverage in mice: the geometry of macronutrient balancing and consequences for fat deposition. Obesity, 16(3), 566–571.

Steele, E. M., Raubenheimer, D., Simpson, S. J., Baraldi, L. G., & Monteiro, C. A. (2018). Ultraprocessed foods, protein leverage and energy intake in the USA. Public health nutrition, 21(1), 114–124.

Tasnin, M. S., Merkel, K., & Clarke, A. R. (2020). Effects of advanced age on olfactory response of male and female Queensland fruit fly, Bactrocera tryoni (Froggatt) (Diptera: Tephritidae). Journal of Insect Physiology, 122, 104024. https://doi.org/https://doi.org/10.1016/j.jinsphys.2020.104024

Terashima, J., & Bownes, M. (2004). Translating Available Food Into the Number of Eggs Laid by Drosophila melanogaster. Genetics, 167(4), 1711–1719. https://doi.org/10.1534/genetics.103.024323

Tsukamoto, Y., Kataoka, H., Nagasawa, H., & Nagata, S. (2014, 2014-March-13). Mating changes the female dietary preference in the two-spotted cricket, Gryllus bimaculatus [Original Research]. Frontiers in Physiology, 5(95), 1–6. https://doi.org/10.3389/fphys.2014.00095

Walker, S. J., Goldschmidt, D., & Ribeiro, C. (2017). Craving for the future: the brain as a nutritional prediction system. Current Opinion in Insect Science, 23, 96–103.

Wysocki, C. J., & Pelchat, M. L. (1993). The effects of aging on the human sense of smell and its relationship to food choice. Critical reviews in food science and nutrition, 33(1), 63–82.

