## Supplementary Figure 1 for "Ageing impairs protein leveraging in a sex-specific manner in *Drosophila melanogaster*"

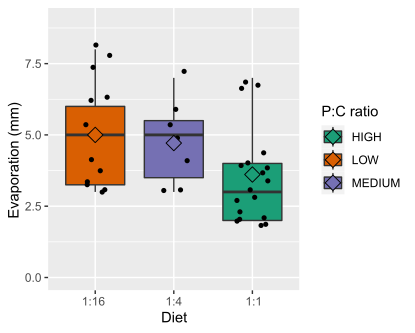

Supplementary figure 1: Evaporation of food from 5 µL capillary tubes with no flies after 2 hours at 80% humidity and 25°C. An ANOVA revealed no significant effect of diet on evaporation, hence evaporation was taken to be constant across assays and diets (F = 2.77, df = 2, p = 0.76).
